# SBT5.2s are the major active extracellular subtilases processing IgG antibody 2F5 in the *Nicotiana benthamiana* apoplast

**DOI:** 10.1101/2024.04.07.588497

**Authors:** Konstantina Beritza, Pierre Buscaill, Shi-Jian Song, Philippe V. Jutras, Jie Huang, Lukas Mach, Suomeng Dong, Renier A. L. van der Hoorn

## Abstract

Plants offer a powerful platform for recombinant protein production but degradation of recombinant proteins by endogenous proteases is causing severe yield losses. Here, we introduce triple knockout lines for SBT5.2, the major active subtilases in the apoplast of agroinfiltrated *Nicotiana benthamiana*. HIV-neutralising IgG antibody 2F5 is no longer cleaved in the apoplast of *sbt5.2* mutants and these mutants accumulate 3-fold more 2F5 upon transient expression but grow normally. Remarkably, however, 2F5 does not accumulate in the apoplast and is not exposed to SBT5.2 when transiently expressed, uncovering an important controversy regarding the subcellular localisation of IgGs in agroinfiltrated plants.

---

Agroinfiltration of *Nicotiana benthamiana* has emerged as a main protein expression platform in plant science and molecular pharming but yields are often hampered by endogenous proteases degrading recombinant proteins. Many IgG antibodies, for instance, are degraded when expressed in *N. benthamiana* (Niemer et al., 2014). HIV neutralising IgG antibody 2F5 is cleaved in the H3 loop of the variable region of the heavy chain (HC) when incubated in apoplastic fluids (AF) of *N. benthamiana*, and this processing was blocked with Ser protease inhibitor PMSF (Niemer et al., 2014). Since the SBT5.2a subtilase (previously called SBT1) is the most abundant active Ser protease in the AF detected by activity-based proteomics (Jutras et al., 2019; Puchol Tarazona et al., 2021), this subtilase was heterologously expressed and shown to cleave 2F5 *in vitro* (Puchol Tarazona et al. 2021). Here, we investigated if SBT5.2 is also necessary to cleave 2F5 in AF.

We first found that fluorescently labelled 2F5 is similarly cleaved as unlabelled 2F5 in AF (Supplemental **Figure S1**). Processing of fluorescent 2F5 is blocked by PMSF and in AF of plants transiently expressing subtilase inhibitor Epi1 but not expressing other protease inhibitors (Supplemental **Figure S2**; Grosse-Holz et al., 2019). Virus-induced Gene Silencing (VIGS) using Tobacco Rattle Virus (TRV) carrying fragments targeting the detected apoplastic subtilases revealed that AF of only *TRV::SBT5.2* plants was unable to cleave 2F5 (Supplemental **Figure S3**), indicating that SBT5.2 is responsible for cleaving 2F5 in AF.

The *SBT5.2* fragment used for VIGS targets three *SBT5.2* homologs that are expressed in agroinfiltrated leaves (Supplemental **Figure S4** and **S5**). Genome editing using CRISPR/Cas9 was used to disrupt all three *SBT5.2* genes, resulting in two independent *sbt5.2* knockout lines that grow indistinguishable from wild-type plants (Supplemental **Figure S6**). Activity-based profiling with FP-TAMRA demonstrated that these lines lack the most active subtilases in the AF, shown at 65-70 kDa in wild-type (WT) plants (**Figure 1a**). AF from *sbt5.2* mutants was unable to cleave 2F5 (**Figure 1b**), demonstrating that SBT5.2s are necessary for 2F5 cleavage.

**Figure 1.**
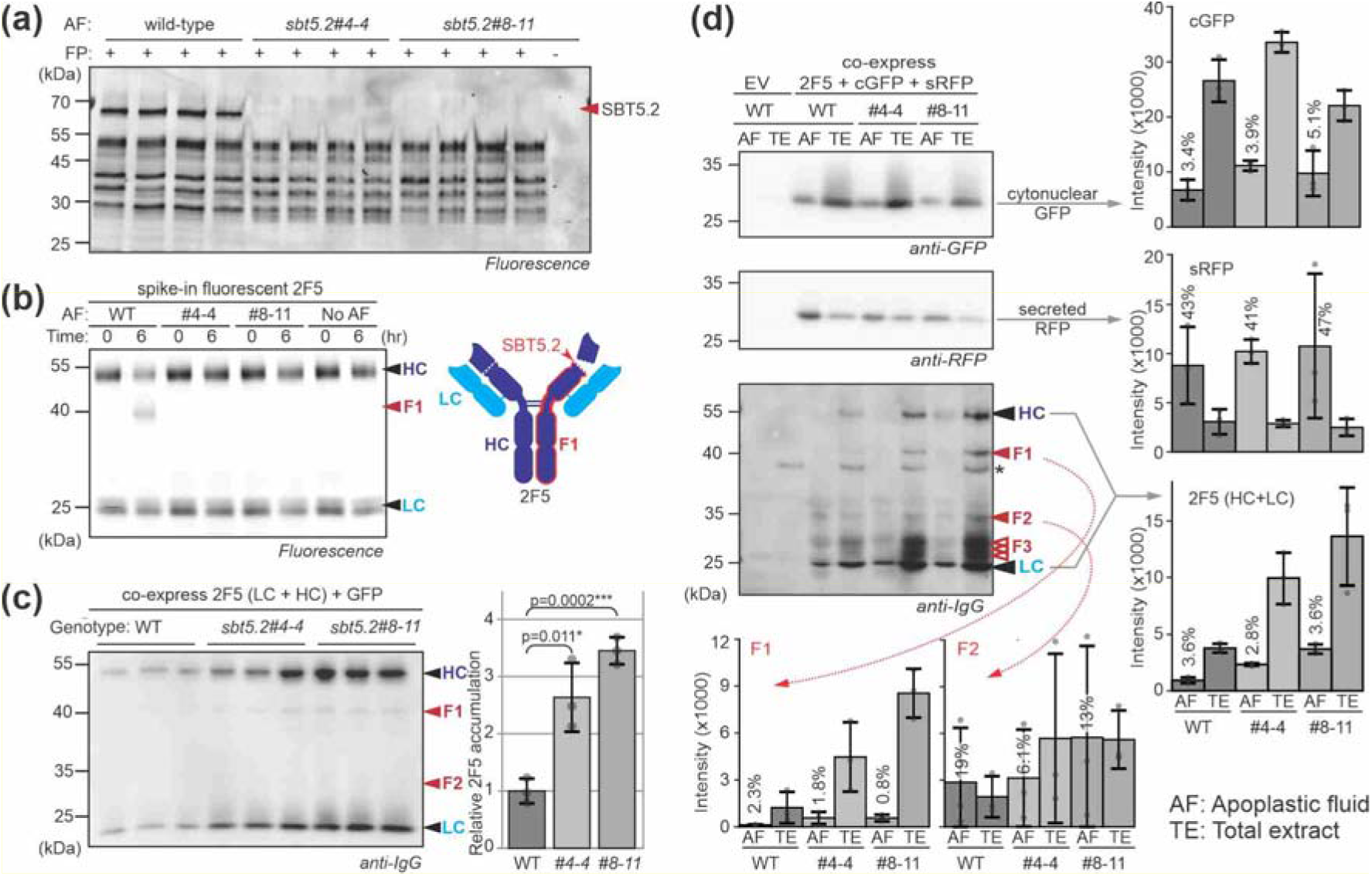
Deletion of three SBT5.2 proteases from *N. benthamiana* avoids 2F5 processing in the apoplast and increases 2F5 accumulation, even though 2F5 is not secreted. **(a)** SBT5.2 are major subtilases in the apoplast. AF was isolated from WT plants and two *sbt5.2* mutants (4 replicates) and active Ser hydrolases were detected with FP-TAMRA and fluorescence scanning. **(b)** 2F5 is no longer cleaved in AF of *sbt5.2* plants. Fluorescently labelled 2F5 was incubated with AFs from WT plants and *sbt5.2* mutants for 0-6 hrs and samples were separated on reducing gels and scanned for fluorescence. **(c)** The *sbt5.2* mutants accumulate 3-fold more 2F5 upon transient expression. HC and LC of 2F5 were transiently co-expressed and signals were quantified from the anti-IgG western blot from a reducing gel. Error bars represent SE of n=3 replicates. **(d)** Transiently expressed 2F5 is not secreted into the apoplast. Both HC and LC of 2F5 were co-expressed with secreted sRFP and cytonuclear cGFP and TEs and AFs were isolated after 5 days from the same leaves in triplicate and analysed by western blotting from a reducing gel using antibodies against GFP, RFP and IgG. Luminescence was quantified and plotted for each signal. F3 fragments are not detected in all experiments. The percentages indicate how much compared to the TE sample was detected in the AF. *, background signal. Error bars represent SE of n=3 replicates.

Transient expression of the HC and light chain (LC) of 2F5 resulted in 3-fold more 2F5 accumulation in total extracts (TEs) of *sbt5.2* mutants compared to WT plants (**Figure 1c**). Also the Ebola-neutralising IgG 2G4 accumulates more in *sbt5.2* mutants (Supplemental **Figure S7**). Fluorescence of co-expressed GFP was similar between WT plants and *sbt5.2* mutants (Supplemental **Figure S8**), indicating that SBT5.2 depletion promotes 2F5 accumulation post-transcriptionally. Remarkably, however, 2F5 processing is unaltered in *sbt5.2* mutants (**Figure 1c**). Similar results were obtained upon co-expression with Epi1 and in *TRV::SBT5.2* plants (Supplemental **Figure S9**), indicating that 2F5 is not exposed to SBT5.2 in the apoplast.

To investigate if 2F5 is secreted in agroinfiltrated plants, we transiently co-expressed 2F5 (HC + LC) with cytonuclear GFP and secreted RFP and isolated AF and TE from the same leaves. Western blot analysis revealed that only 3-5% of the GFP detected in TE is detected in AF whereas 41-47% of the total RFP is in the AF (**Figure 1d**). Neither GFP nor RFP distribute differently nor accumulate higher in the *sbt5.2* mutants. Importantly, only 2-4% of both HC and LC of 2F5 were detected in AF (**Figure 1d**), indicating that 2F5 is not secreted and not exposed to SBT5.2 in the apoplast. Fragment F1 follows the same trend as HC and LC signals, while fragment F2 accumulates relatively abundantly in AF (**Figure 1d**). F2 is also easily detected in AF (Niemer et al., 2014) and originates from degradation of unassembled HC because it also accumulates when only HC is expressed (Supplemental **Figure S10**). The distribution of 2F5 signals is similar between WT and *sbt5.2* plants but 3-fold more 2F5 accumulates in *sbt5.2* mutants (**Figure 1d**), consistent with **Figure 1c**.

The observation that most 2F5 is not secreted challenges the general assumption that IgGs are secreted and seems to contradict numerous reports on the detection of IgGs in the apoplast (e.g. Arcalis et al., 2013; Ocampo et al., 2016). Our data is nevertheless consistent with the literature. A previous study on transient 2F5 expression revealed that only an estimated 5% of the HC signal detected in TE was detected in AF (Niemer et al., 2014). Fluorescent fusion proteins of the HC of 2F5 were found to accumulate in pre-vacuolar compartments and the vacuole (Irons et al., 2008). Similar observations were made when studying 2G12, a different HIV-neutralising IgG (Irons et al., 2008), and 2G4 (Supplemental **Figure S7**). It has been speculated that secretion is prevented either by cryptic vacuolar targeting signals or by chaperone BiP, which remains bound to IgGs (Irons et al., 2008). Thus, besides introducing a very useful protease mutant to the community to increase yields, this study also highlights an important controversy on the subcellular accumulation of IgGs in agroinfiltrated plants.

## Acknowledgements

We thank Urszula Pyzio for plant care and Sarah Rodgers, Caroline O’Brian and Patricia Bowman for technical support.

## Author contributions

RH conceived and managed the project; KB performed the majority of the experiments; PB produced VIGS constructs, initiated silencing experiments and conducted phylogenetic analysis; PJ produced pPJ057 (sRFP); SS selected *sbt5.2* lines with help from JH and SD; LM assisted with data interpretation; KB and RH wrote the article with input from all co-authors.

## Data availability

All data are provided as figures and supplemental figures.

