## Supplemental for "SBT5.2s are the major active extracellular subtilases processing IgG antibody 2F5 in the *Nicotiana benthamiana* apoplast"

#### Supplemental Materials & Methods

**Bacterial strains and growth conditions -** The *Escherichia coli* strain used for plasmid propagation was DH10β on lysogeny broth (LB) or agar, supplemented with appropriate antibiotics. Following heat shock transformation, *E. coli* transformants were selected on LB agar medium plates containing 50 μM kanamycin or carbenicillin. The *Agrobacterium tumefaciens* strain GV3101-pMP90 was used for transient transformation of *Nicotiana benthamiana*. Following heat shock transformation, *A. tumefaciens* transformants were selected on LB agar medium plates containing 25 μM rifampicin, 10 μM gentamycin, and 50 μM kanamycin or carbenicillin for the binary vector.

***Nicotiana benthamiana* cultivation -** Wild-type *N. benthamiana* (LAB strain; Ranawaka et al., 2023) seeds were sown into a 3:1 mix of soil (Sinclair Modular Seed Peat reduced propagation mix) with vermiculite (Sinclair Pro Medium) in 7 cm square pots and grown at high humidity under transparent plastic covers for 5 days. Seedlings were uncovered and grown first for one week in the greenhouse at 22-23°C, 16 hr light/8 hr dark, at high humidity, and 80-120μmol/$m^2$s light intensity and watered three times per week.

**Molecular cloning -** The HC and LC of 2F5 (PDB: 2p8l) and 2G4 were codon optimised for *N. benthamiana* and obtained from Twist Bioscience, carrying BsaI sites for Golden Gate cloning (Supplemental **Table S1**, Engler et al., 2014; Marillonnet & Werner, 2015; Werner et al., 2012). All overexpression constructs carry the N-terminal PR1a signal peptide from *N. tabacum* unless stated otherwise (Supplemental **Table S2**). All constructs were cloned into a Golden Gate binary vector containing the T-DNA with a Cauliflower Mosaic Virus 35S (CaMV 35S) promoter and either a nopaline synthase (nos) or 35S terminator following the Golden Gate protocol in a ratio of insert:backbone 2:1. Similarly, for designing the VIGS plasmids, the *Nb*SBTs genes were selected by a BLAST search in the Sol genomics network database (https://solgenomics.net/). The gene fragments (Supplemental **Table S3**) targeting the subtilisin-like proteases were obtained from Invitrogen and were inserted into pJK037 (Morimoto et al., 2022) using Golden Gate cloning (BsaI sites), to generate the *TRV-GFP* and *TRV-NbSBT* plasmids (Supplemental **Table S2**). The protease inhibitor constructs were previously cloned and described (Grosse-Holz et al., 2018b). RFP was amplified from secRFP (Samalova et al., 2006) using primers 5'-ttgtggtctcaaatgaagactaatctttttctctttctcatcttttcac-3' and 5'-ttcgtggtctcaaagcctaggcgccggtggag-3' and inserted into pJK268c (Kourelis et al., 2020) using Golden Gate cloning to produce binary plasmid pPJ057, which carries a T-DNA with silencing suppressor p19 and an 2x35S-driven secreted RFP.

**Agroinfiltration of *Nicotiana benthamiana* -** *Agrobacterium* cultures were resuspended in infiltration buffer (10 mM $MgCl_2$, 10 mM MES-K/ pH 5.7 with KOH, with filter-sterilized 100 μM acetosyringone added immediately before use) to reach a final $OD_{600}$ of 0.3-0.5. Leaves of approximately 4-week-old *N. benthamiana* were used for infiltration, while the same leaf was infiltrated for each technical replicate to achieve the same developmental stage. After infiltration, the plants were grown for 5 days post-infiltration (dpi) in a growth chamber at 22-23 °C, 16 hr light/ 8hr dark, at high humidity and 80-120 μmol/$m^2$s light intensity, and 1 cm diameter leaf discs were harvested for analyses. For *in vivo* experiments six 1 cm diameter leaf discs were pooled per technical replicate, resulting in six biological replicates from three individual technical replicates (n=18).

**Virus-induced gene silencing -** The TRV1 and TRV2 plasmids were transformed into *Agrobacterium* strain GV3101 and mixed in a 1:1 ratio to a final $OD_{600}$ of 0.5 for agroinfiltration. Two weeks old *N. benthamiana* were agroinfiltrated with *TRV::NbSBT* or *TRV::GFP*, and 3 weeks after agroinfiltration successful silencing was assessed by checking for bleached leaves in *TRV::PDS* plants (Supplemental **Figure S7**).

**Generation of *sbt5.2* triple mutants** – *N. benthamiana* LAB was transformed with a T-DNA carrying Cas9 and two single guide RNAs targeting each gene (Supplemental **Table S4**). The *sbt5.2* triple mutants were generated using CRISPR/Cas9 editing by Biogle (Hangzhou, China). Primary transformants were selected for carrying mutations in the target genes by sequencing PCR products generated from genomic DNA using sequencing primers listed in Supplemental **Table S4**. T2 plants were screened for homozygosity of the mutant alleles using the same sequencing primers.

**Total and apoplastic fluid extraction -** All samples were harvested 5 days post infiltration. For total extracts, six leaf discs (1 cm diameter) were harvested and ground to a fine powder using liquid nitrogen, and 150 µl phosphate-buffered saline (PBS) supplemented with Halt™ Protease Inhibitor Cocktail (Thermo 78430) was added. Samples were then centrifuged for 25 minutes at 4°C and 13,000 x *g* and the supernatant was transferred into a new tube with loading dye for protein gel loading. For apoplastic fluids, the respective six leaves following leaf disc harvest were vacuum infiltrated with Milli-Q water. The leaf surfaces were then dried with tissue paper, and centrifuged in a 25 mL syringe hanging in a 50 mL tube at 1000 x *g* for 25 minutes at 4 °C. Aliquots of apoplastic fluids were collected from the 50 mL tube and were flash-frozen and stored at -20 °C until further use.

**Antibody labelling -** The 2F5 antibody was provided by Polynum Scientific (anti-gp41; AB001). 2F5 was labelled using an amine-reactive dye (DyLight® 488 NHS Ester; Thermo 46402) by diluting 0.06 mg 2F5 in 100 µl PBS (pH 7.4) and adding 0.2 µg dye. The mix was incubated for 1 hour in the dark at room temperature. Excess dye was removed by loading the labelling mix into a column (Amicon® Ultra 0.5 centrifugal filter devices 3K; Merck UFC500324) and centrifuged for 8 minutes at 10,000 x *g*. After loading another 100 µl PBS, the column was centrifuged for 8 minutes at 10,000 x *g*. The above step was repeated, the column was transferred upside-down into a new tube and centrifuged for 5 minutes. The labelled antibody was kept in aluminium foil at 2-8 °C and used within a month.

***In vitro* degradation assays -** Fluorescently labelled 2F5 antibody was incubated in apoplastic fluid from *N. benthamiana* leaves. Samples were taken at zero and six-hour time points. The reaction was stopped by adding 4x loading dye with dithiothreitol (0.2 M Tris pH 6.8, 8% w/v SDS, 40% w/v glycerol, 0.4% w/v bromophenol blue, 0.6 M DTT; Thermo R0862). The samples were then incubated at 95 °C for 5 minutes before sodium dodecyl-sulfate polyacrylamide gel electrophoresis (SDS-PAGE). The effect of protease inhibitors on the degradation of 2F5 in wild-type apoplastic fluid was investigated by pre-incubation for 30 minutes using chemical protease inhibitors 100 µM E-64 (Merck E3132) or 1 mM phenylmethylsulfonyl fluoride (PMSF; Merck 10837091001). For negative controls, the samples were treated with the same volume of dimethyl sulfoxide (DMSO; Merck D8418).

**SDS-PAGE and western blotting -** The samples were incubated at 95 °C for 5 minutes before loading and separated at 170V in Invitrogen Novex vertical gel tanks. The Amersham® Typhoon (Cy2: 488 nm for labelled antibody, Cy5: 685 nm for ladder) was used to measure fluorescence. The proteins were also visualised by total protein staining with Instant Blue® Coomassie Protein Stain (Abcam ab119211) or were transferred onto a polyvinylidene difluoride (PVDF, BioRad) membrane for western blotting. For Western blot analysis, proteins were transferred to PVDF membranes using BioRad Trans-blot Turbo® according to the manufacturer's instructions (BioRad Kit 1704275). Blots were blocked for 2 hours or overnight with 5% (w/v) skim milk in PBS-T (PBS tablets; Merck 524650, 0.1% Tween-20; Merck P1379). For IgG antibody detection, the membrane was incubated with 1:3000 HRP-conjugated anti-human IgG in 5% (w/v) skim milk for 2 hours at room temperature. Blots were washed twice with PBS-T for 5 minutes, developed using SuperSignal™ West Pico PLUS Chemiluminescent Substrate (Thermo 34580), and visualised using the ImageQuant® LAS-4000 imager. Image data were quantified using the open-source software ImageJ, and background values were subtracted based on control images.

**Activity-based protein profiling (ABPP) -** ABPP was conducted for the *sbt5.2* knockout plants, using the fluorophosphonate (FP) TAMRA probe for targeting Serine hydrolases as described previously (Jutras et al., 2019). Briefly, 48 µL of apoplastic fluids, were adjusted to a solution containing 50 mM sodium acetate (NaAc) at pH 5 and 5 mM dithiothreitol (DTT). The mixture was then incubated for 1 hour with 0.5 µM FP-TAMRA (Thermo 88318). To stop activity-based labelling, 1 mL of cold acetone was added. The samples were then centrifuged, the supernatant was removed, and the proteins were resuspended in a gel-loading buffer. These protein samples were heated and separated on a 12% SDS-PAGE at reducing conditions. SDS-PAGE gels were scanned for in-gel fluorescence using a Typhoon scanner with Cy3: 550 nm settings. Four biological replicates were used in each case for analysis (n=4).

**Estimating secretion efficiency -** Total leaf extracts (TE) samples derived from six individual leaf discs pooled together and mixed with 150 µl PBS buffer supplemented with protease inhibitor cocktail. Apoplastic fluids (AF) samples derived from pooling the same six leaves, following leaf disc removal used for TE, leading to 400 µl sample volume. The leaf areas were calculated using ImageJ (Polygon Tool and Analyze>Measure), converted to Total Leaf Area ($cm^2$) and normalised for the Loaded Fraction which represents the sample used for SDS-PAGE loading as follows:

$$\textit{Total Leaf Area (AF)} = \frac{\sum whole\ leaf\ area - 6\ x\ leaf\ disc\ area}{area\ of\ 1cm^2}$$

$$\textit{Total Leaf Area (TE)} = \frac{6\ x\ leaf\ disc\ area}{area\ of\ 1cm^2}$$

$$\textit{Loaded Fraction (AF or TE)} = \frac{Total\ Leaf\ Area\ (AF\ or\ TE)}{percentage\ of\ loaded\ sample\ (\%;\ AF\ or\ TE)}$$

Western blot band intensities were calculated in ImageJ (Analyze>Gels>Plot Lanes) by calculating the area of the grey value distribution. The raw intensities were normalised to correspond to the respective normalised Loaded Fraction as follows:

$$\textit{Normalised intensities (AF or TE)} = \frac{Raw\ intensity\ (AF\ or\ TE)}{Loaded\ Fraction\ (AF\ or\ TE)}$$

Total leaf extracts contain the total protein content found in samples, so a secreted protein will appear in both the TE and the AF samples, with most of the protein being represented in AF samples as follows:

$$\textit{Percentage of secretion (\%)} = \frac{Normalised\ intensities\ (AF)}{Normalised\ intensities\ (TE)}\ x\ 100$$

**Phylogenetic analysis by Maximum Likelihood -** The proteomes of *N. benthamiana* and *Arabidopsis thaliana* were retrieved from the NbDE database (Kourelis et al. 2019), and the Arabidopsis Information Resource (TAIRv10.1), respectively. The proteomes of *Oryza sativa, Physcomitrium patens, Solanum lycopersicum,* and *Zea mays* were obtained from GenBank (National Center for Biotechnology Information [NCBI]). These proteomes were annotated with PFAM identifiers (El-Gebali et al., 2019), and all sequences containing a peptidase S8 (PF00082) domain were extracted from the different datasets.

The predicted amino acid sequences were aligned using Clustal Omega (Lemoine et al., 2019). Alignment curation involved trimming the sequences with TrimAL (Capella-Gutiérrez et al., 2009; Lemoine et al., 2019). Additionally, partial and truncated S8 proteins, as well as S8 sequences with an incomplete catalytic triad, were manually removed. When isoforms resulted in the same S8 domain, only one sequence was conserved. Isoforms are indicated as follows: At1g20160.1, .2, .3.

The evolutionary analysis was conducted using the Maximum Likelihood method and the Whelan And Goldman model (Whelan and Goldman, 2001) in MEGA X (Kumar et al., 2018). The bootstrap consensus tree inferred from 500 replicates (Felsenstein 1985) represents the evolutionary history of the taxa analysed. Branches corresponding to partitions reproduced in less than 50% bootstrap replicates are collapsed. The percentage of replicate trees in which the associated taxa clustered together in the bootstrap test (500 replicates) is shown next to the branches. The initial tree for the heuristic search were obtained automatically by applying Neighbor-Join and BioNJ algorithms to a matrix of pairwise distances estimated using the JTT model. The topology with the superior log-likelihood value was selected. A discrete Gamma distribution was used to model evolutionary rate differences among sites (5 categories (+G), parameter = 1.0885). This analysis involved 346 amino acid S8 domain sequences. All positions with less than 80% site coverage were eliminated, i.e., fewer than 20% alignment gaps, missing data, and ambiguous bases were allowed at any position (partial deletion option). A total of 392 sequences were in the final dataset. The S8 domain (130-374 aa) from *Bacillus subtilis* subtilisin (O87655) was used to root the phylogenetic trees.

The naming of subtilase clades (SBT1-6) follows (Rautengarten et al., 2005). The naming of *N. benthamiana* subtilases (NbSBT) is based on the percentage identity to the closest Arabidopsis ortholog. For instance, NbD038072, NbD021558, and NbD013006 have 57.70%, 55.75%, and 54.51% identity, respectively, to AtSBT5.2 and consequently are named NbSBT5.2a, NbSBT5.2b, and NbSBT5.2c, respectively.

### SUPPLEMENTAL FIGURES

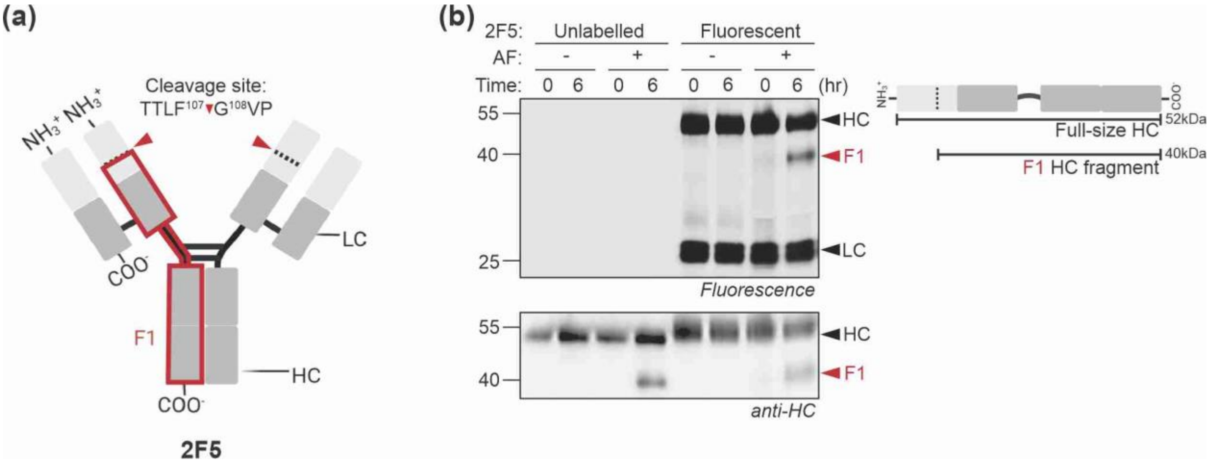


**Figure S1** Amine-reactive dye labelling does not affect 2F5 cleavage *in vitro*.

**(a)** Schematic representation of IgG antibody 2F5 with two LC and two HC, with the red arrow indicating the previously described cleavage site (Puchol-Tarazona et al., 2021), caused by cleavage of 2F5 in the variable region of the HC, resulting in 12 and 40 kDa fragments. The detected 40 kDa F1 fragment is highlighted with a red box. **(b)** Processing of labelled and unlabelled 2F5 in apoplastic fluids. 2F5 was incubated with and without apoplastic fluids for 0 or 6 hours, separated on reducing protein gels and analysed by fluorescence scanning and western blotting using anti-HC antibodies.

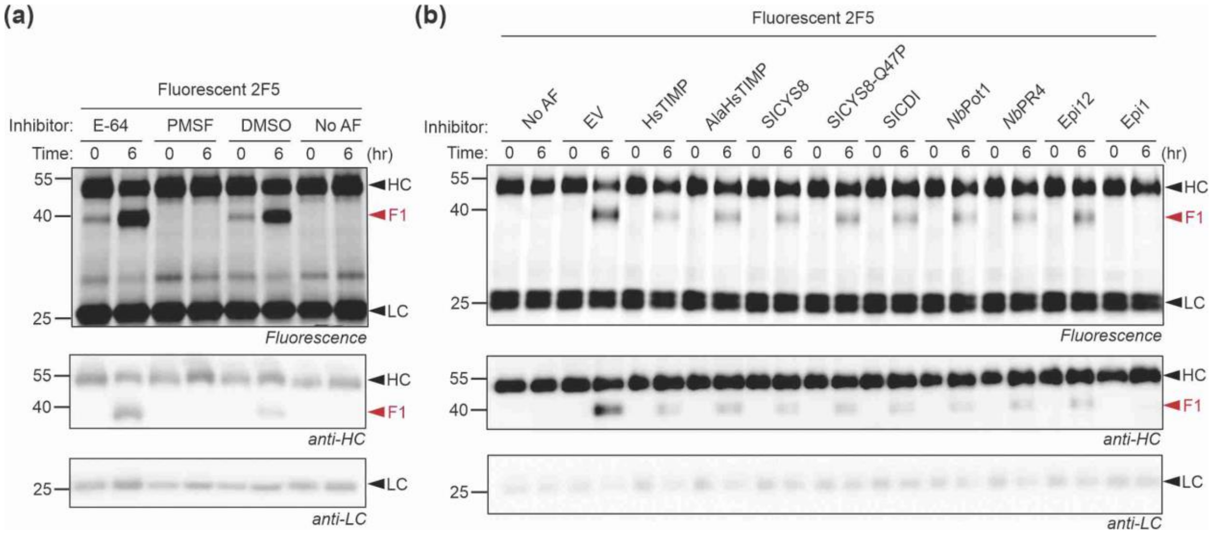


**Figure S2** PMSF and Epi1 block 2F5 processing in apoplastic fluids.

**(a)** PMSF blocks 2F5 processing. Apoplastic fluid or water was incubated with Cys protease inhibitor E-64, Ser protease PMSF or DMSO (negative control) and 2F5 was added for 0 or 6 hours, separated on reducing protein gels and analysed by fluorescence scanning and western blotting using anti-HC and anti-LC antibodies. **(b)** Epi1 blocks 2F5 processing. Apoplastic fluids from plants transiently expressing various inhibitors were incubated with fluorescent 2F5 for 0 or 6 hours, separated on reducing protein gels and analysed by fluorescence scanning and western blotting using anti-HC and anti-LC antibodies. Apoplastic fluids from empty vector (EV) agroinfiltrated plants and water were used as positive and negative controls, respectively.

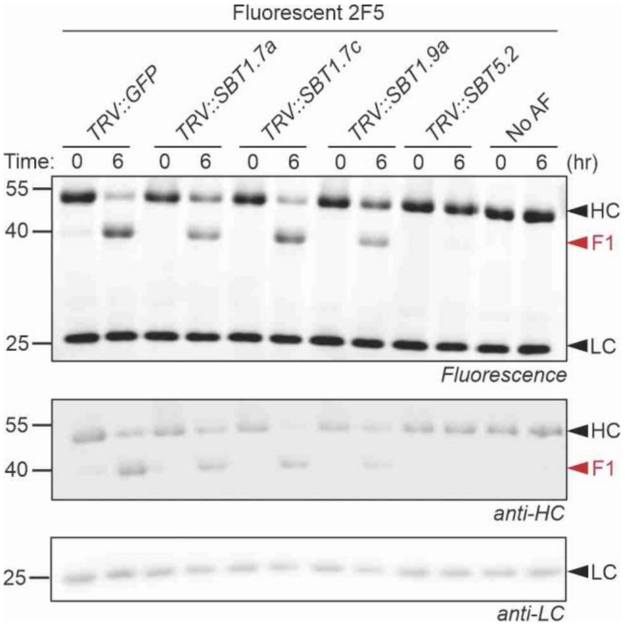


**Figure S3** *SBT5.2* silencing prevents 2F5 processing in apoplastic fluids.
Apoplastic fluids from plants silenced for different subtilases were incubated with fluorescent 2F5 for 0 or 6 hours, separated on reducing protein gels and analysed by fluorescence scanning and western blotting using anti-HC and anti-LC antibodies.

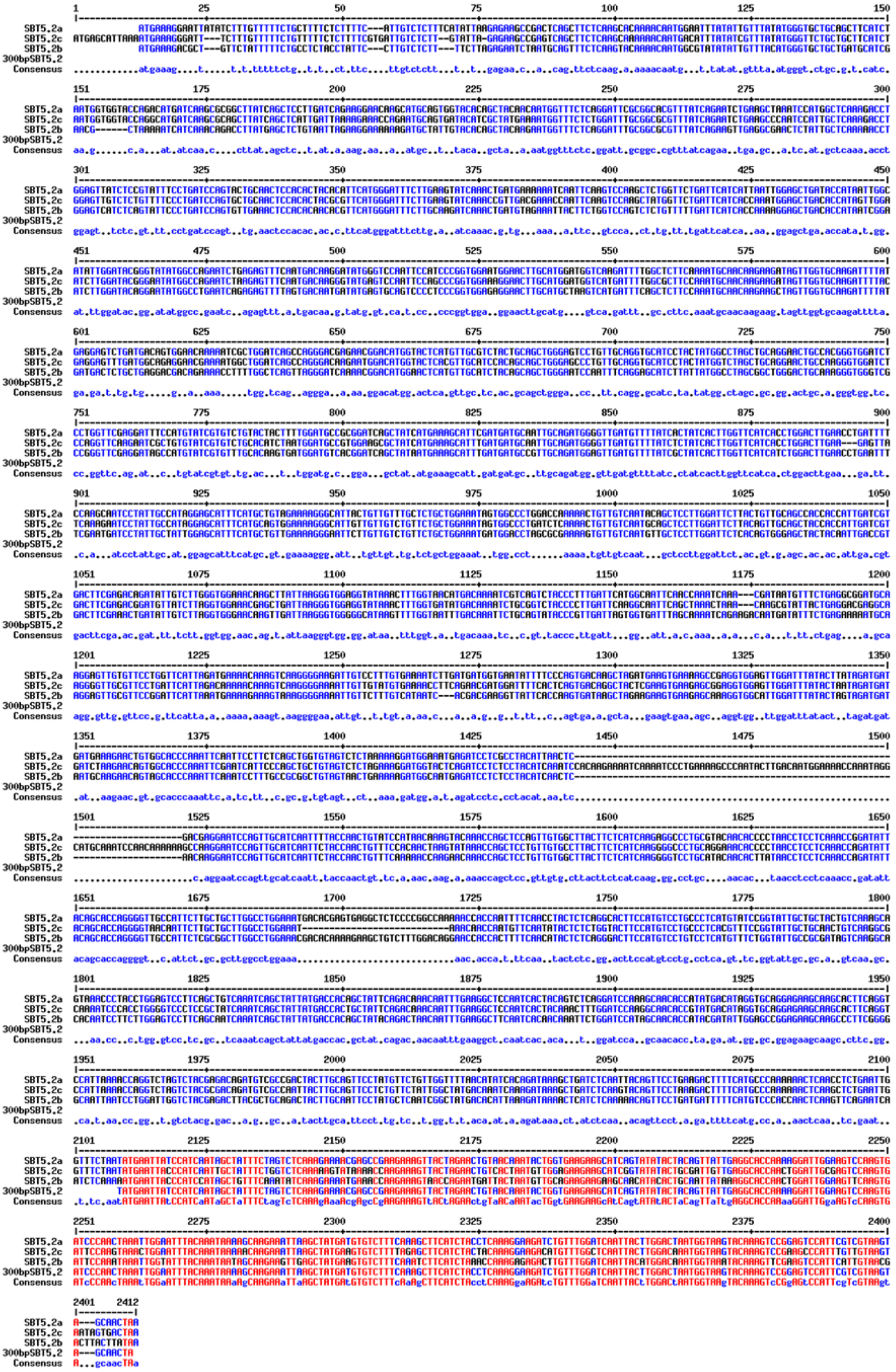

**Figure S4** VIGS targets three *SBT5.2* genes.

The 300bp fragment used for *SBT5.2* silencing has sufficient homology to silence all three SBT5.2-encoding genes.

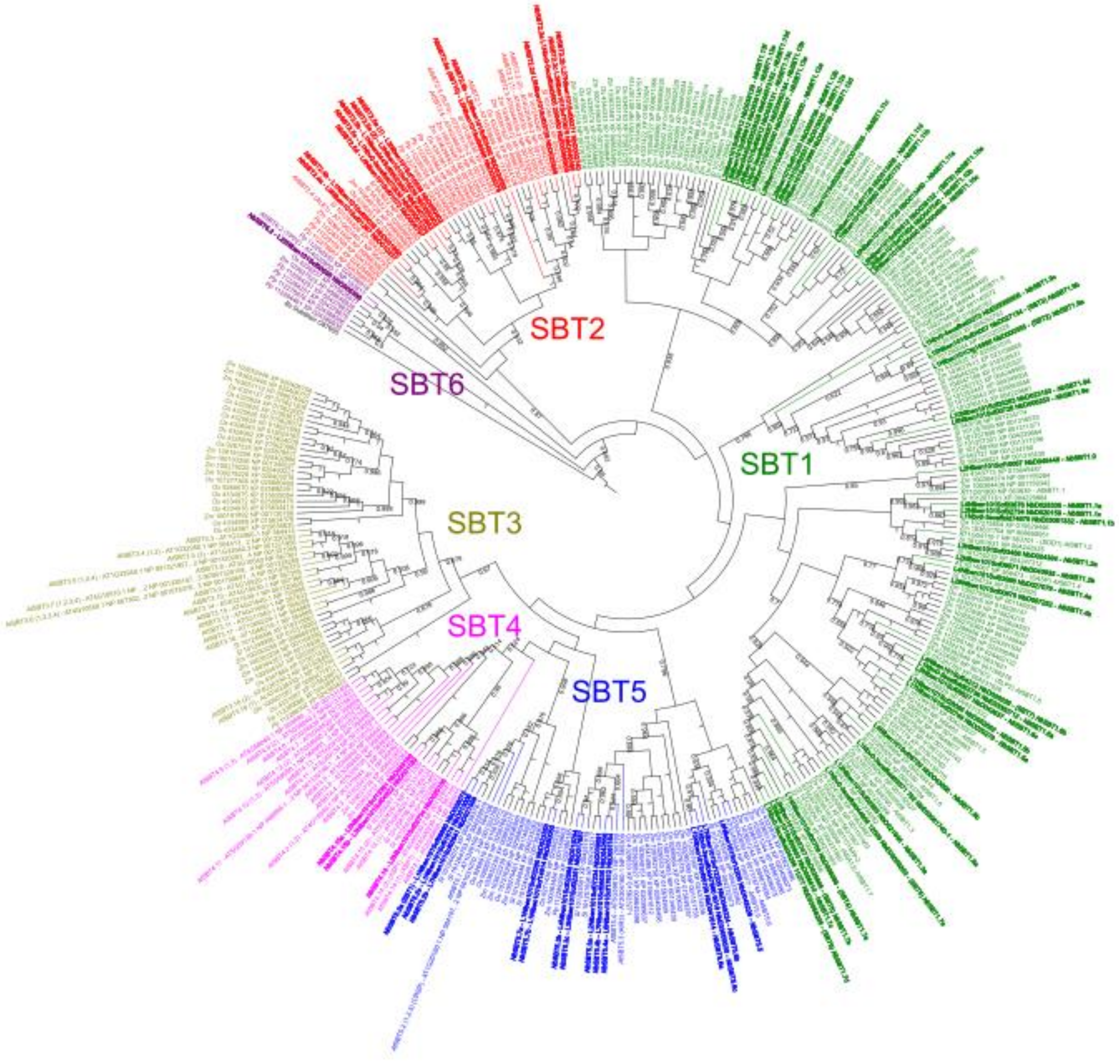


| NbDE | Lab360 | Final name | Mock | | Agroinfiltrated | |
|---|---|---|---|---|---|---|
| | | | mean | SD | mean | SD |
| NbD038072 | NbL13g04590 | *Nb*SBT5.2a | 0 | 0 | 0.093467031 | 0.042122779 |
| NbD021558 | NbL04g13700 | *Nb*SBT5.2b | 9.744890966 | 2.14370341 | 4.553243162 | 0.921318851 |
| NbD013006 | NbL13g04600 | *Nb*SBT5.2c | 0.618187564 | 0.27390427 | 0.523724281 | 0.303865567 |

**Figure S5** Phylogenetic analysis of subtilases and RPKM values for three *NbSBT5.2* genes. **Top:** Phylogenetic analysis of the catalytic domain of subtilases from *N. benthamiana (Nb*, **bold**), *Arabidopsis thaliana (At)*, *Oryza sativa (Os)*, *Physcomitrium patens (Pp)*, *Solanum lycopersicum (Sl)*, and *Zea mays (Zm)*. Subtilases form six subfamilies, SBT1-6**. Bottom:** Reads Per Kilobase of transcript per Million mapped reads (RPKM) values of *Nb*SBT5.2a, *Nb*SBT5.2b, and *Nb*SBT5.2c in mock or agroinfiltrated plants (Grosse-Holz et al., 2018a)**.**

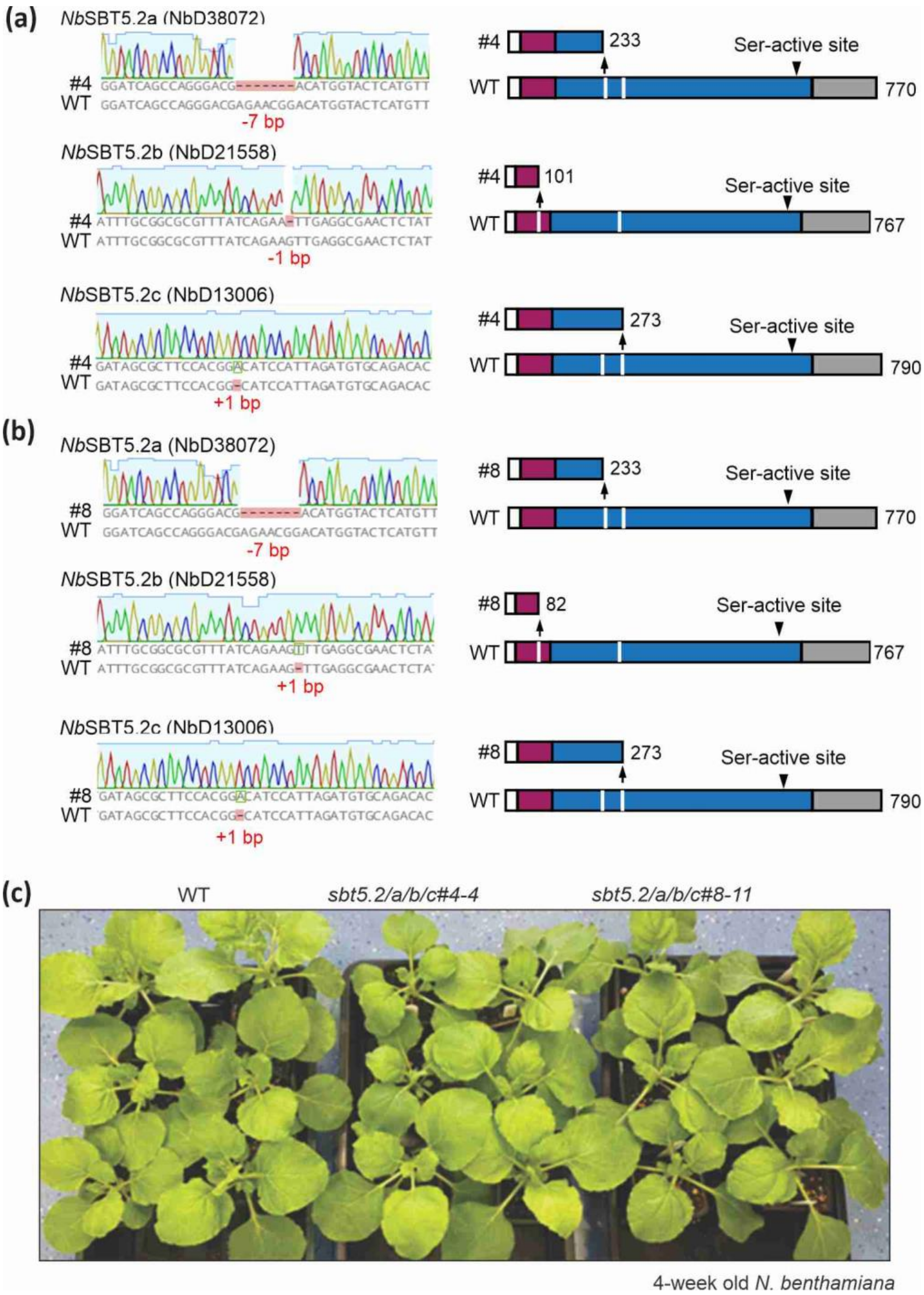


**Figure S6** Identification of *sbt5.2* triple KO mutants in *N.benthamiana*.
**(a) Left:** Sequencing results of NbSBT5.2a, NbSBT5.2b and NbSBT5.2c obtained from two CRISPR-Cas9-generated lines *sbt5.2* #4 and *sbt5.2* #8. **Right:** The predicted protein domains of *Nb*SBT5.2a, *Nb*SBT5.2b, and *Nb*SBT5.2c in the mutant plant (top bar) compared to the wild-type lines (bottom bar). Signal peptides are represented by white boxes, the inhibitor region is indicated by the magenta box, the subtilase domain is highlighted in blue, and the location of sgRNAs is denoted by white lines.

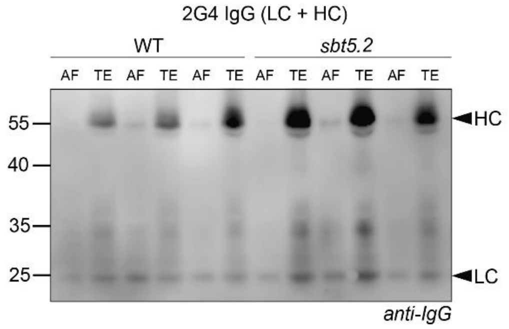


**Figure S7** IgG 2G4 also accumulates more in *sbt5.2* mutant but not in the apoplast.
The HC and LC of 2G4 were co-expressed by agroinfiltration and apoplastic fluids (AF) and total extracts (TE) were harvested 5 days later in triplicate and analysed by western blot using anti-IgG antibodies.

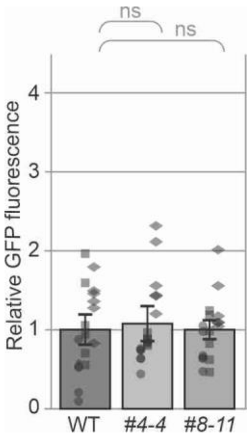

**Figure S8** Transiently co-expressed GFP is not expressed more in the sbt5.2 mutants. Cytonuclear GFP was transiently co-expressed with 2F5 and GFP fluorescence was quantified at day 5 from six different leaves in three experimental replicates. Error bars represent SE of n=3 experimental replicates.

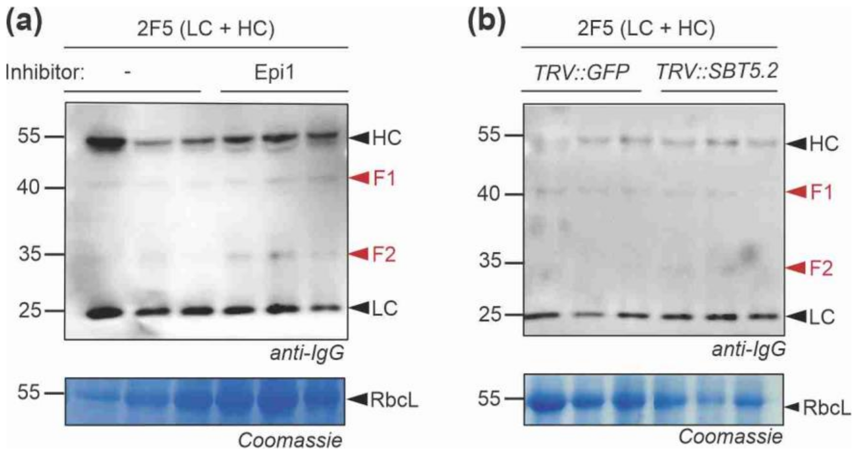


**Figure S9** SBT5.2 depletion with Epi1 or VIGS does not affect 2F5 processing *in vivo*

**(a)** Epi1 has no impact on processing of transiently expressed 2F5 *in vivo*. HC and LC of 2F5 were transiently co-expressed with and without Epi1 for 5 days by agroinfiltration in triplicate. Total extracts were generated and separated by reducing SDS-PAGE and analysed by western blot using anti-HC and -LC antibodies. F1 and F2 indicate two different fragments of the 2F5 HC. **(b)** *SBT5.2* silencing has no impact on the processing of transiently expressed 2F5 *in vivo*. HC and LC of 2F5 were transiently co-expressed in *TRV::GFP* and *TRV::SBT5.2* plants for 5 days by agroinfiltration in triplicate.

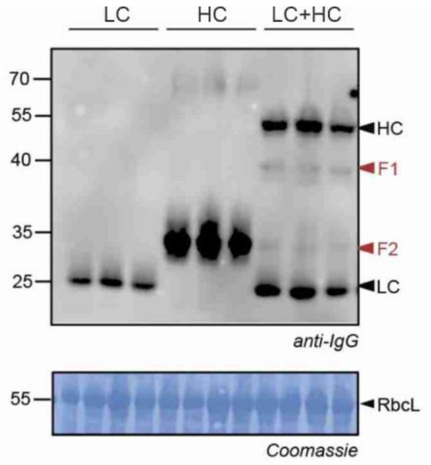


**Figure S10** Fragment F2 originates from HC overexpression.

The LC, HC and LC+HC of 2F5 were transiently expressed by agroinfiltration in *N. benthamiana* and total extracts were generated five days later in triplicate and analysed by Coomassie staining and western blotting using anti-IgG antibody.

**Table S1** Coding sequences for 2F5 and 2G4 expression

| |
|---|
| >PR1a_2F5_LC_CDS<br>ATGGGATTTGTTCTCTTTTCACAATTGCCTTCATTTCTTCTTGTCTCTACACTTCTCTTATTCCTAGTAATATCCCACTCTTGCCGTGCAG<br>GTGCATTACAATTAACACAATCTCCCTCCTCACTTAGTGCCTCCGTGGGGGATCGCATAACAATAACATGTAGAGCATCTCAGGGAGTGAC<br>CTCCGCACTTGCATGGTATCGACAAAAGCCAGGATCCCCCCCTCAGTTGTTAATTTATGACGCATCAAGTTTGGAGTCAGGCGTGCCTTCC<br>AGATTTTCTGGTTCCGGTTCAGGAACTGAATTTACACTGACAATAAGTACTCTAAGGCCAGAGGACTTTGCCACTTATTACTGTCAACAAC<br>TACATTTTTACCCCCACACGTTCGGAGGTGGAACAAGGGTCGATGTACGTAGGACTGTTGCTGCTCCATCTGTTTTCATCTTCCCACCATC<br>TGATGAGCAGCTCAAGTCTGGAACTGCTTCTGTTGTTTGCCTCCTCAACAATTTCTACCCAAGGGAAGCTAAGGTGCAGTGGAAAGTTGAT<br>AATGCTCTCCAGTCCGGAAACTCCCAAGAATCTGTTACTGAGCAGGATTCCAAGGATTCTACTTACTCCCTCTCCTCAACTCTCACTCTCT<br>CTAAGGCTGATTACGAGAAGCACAAGGTTTACGCTTGCGAAGTTACTCACCAGGGACTTTCTTCACCAGTGACTAAGTCTTTCAATAGGGG<br>CGAGTGCTAA |
| >PR1a_2F5_LC_aa<br>MGFVLFSQLPSFLLVSTLLLFLVISHSCRAGALQLTQSPSSLSASVGDRITITCRASQGVTSALAWYRQKPGSPPQLLIYDASSLESGVPS<br>RFSGSGSGTEFTLTISTLRPEDFATYYCQQLHFYPHTFGGGTRVDVRRTVAAPSVFIFPPSDEQLKSGTASVVCLLNNFYPREAKVQWKVD<br>NALQSGNSQESVTEQDSKDSTYSLSSTLTLSKADYEKHKVYACEVTHQGLSSPVTKSFNRGEC |
| >PR1a_2F5_HC_CDS<br>ATGGGATTTGTTCTCTTTTCACAATTGCCTTCATTTCTTCTTGTCTCTACACTTCTCTTATTCCTAGTAATATCCCACTCTTGCCGTGCAG<br>GTAGGATCACCCTGAAAGAATCTGGACCTCCTTTGGTGAAGCCTACCCAGACTCTTACTCTGACCTGCTCCTTCTCTGGTTTCTCCCTGTC<br>TGATTTCGGTGTTGGTGTGGGTTGGATTAGACAGCCTCCTGGTAAGGCTCTTGAGTGGCTTGCTATCATCTACTCCGATGATGATAAGAGG<br>TACAGCCCAAGCCTTAACACCAGGCTTACCATCACCAAGGATACCAGCAAGAACCAGGTGGTGCTTGTGATGACTAGGGTTTCACCTGTCG<br>ACACCGCTACCTACTTTTGCGCTCATAGAAGGGGTCCTACCACCCTTTTCGGAGTTCCTATTGCTAGGGGTCCTGTGAACGCTATGGATGT<br>TTGGGGTCAGGGTATTACCGTGACCATCAGCTCTACTTCAACTAAGGGACCATCTGTTTTTCCACTCGCTCCATCCTCTAAGTCTACTTCA<br>GGTGGAACTGCTGCTCTTGGATGCCTTGTTAAGGATTACTTTCCAGAGCCAGTGACTGTGTCTTGGAATTCTGGTGCTCTTACTTCCGGTG<br>TTCATACTTTCCCAGCTGTGCTTCAATCTTCCGGACTTTACTCTCTTTCCTCTGTTGTGACTGTGCCATCTTCTTCACTTGGCACTCAAAC<br>TTACATCTGCAACGTGAACCACAAGCCATCCAACACAAAAGTGGATAAGAAGGTTGAGCCAAAGTCCTGCGATAAGACTCATACTTGTCCA<br>CCATGTCCAGCTCCAGAACTTCTTGGTGGTCCTTCTGTTTTTTTGTTCCCACCAAAGCCAAAGGATACTCTCATGATCTCTAGGACTCCAG<br>AGGTTACATGCGTTGTGGTTGATGTGTCTCATGAAGATCCAGAGGTGAAGTTCAACTGGTATGTGGATGGTGTTGAGGTGCACAACGCTAA<br>GACTAAGCCAAGAGAGGAACAGTACAACTCCACTTACAGGGTTGTGTCTGTGCTTACTGTTCTTCACCAGGATTGGCTTAACGGCAAAGAG<br>TACAAGTGCAAGGTGTCCAACAAGGCTTTGCCAGCTCCAATCGAAAAGACTATCTCTAAGGCTAAGGGACAGCCAAGGGAACCTCAAGTTT<br>ACACTCTTCCACCATCTAGGGATGAGCTTACTAAGAACCAGGTGTCCCTTACTTGCCTTGTGAAGGGATTTTACCCATCCGATATTGCTGT<br>TGAGTGGGAGTCTAATGGACAGCCTGAGAACAACTACAAGACTACTCCACCAGTGCTCGATTCCGATGGATCATTCTTCTTGTACTCCAAG<br>CTCACTGTGGATAAGTCTAGGTGGCAACAGGGAAACGTTTTCTCTTGCTCTGTTATGCATGAGGCTCTCCACAATCACTACACTCAGAAGT<br>CCCTTTCTTTGTCCCCTGGCAAGTAA |
| >PR1a_2F5_HC_aa<br>MGFVLFSQLPSFLLVSTLLLFLVISHSCRAGRITLKESGPPLVKPTQTLTLTCSFSGFSLSDFGVGVGWIRQPPGKALEWLAIIYSDDDKR<br>YSPSLNTRLTITKDTSKNQVVLVMTRVSPVDTATYFCAHRRGPTTLFGVPIARGPVNAMDVWGQGITVTISSTSTKGPSVFPLAPSSKSTS<br>GGTAALGCLVKDYFPEPVTVSWNSGALTSGVHTFPAVLQSSGLYSLSSVVTVPSSSLGTQTYICNVNHKPSNTKVDKKVEPKSCDKTHTCP<br>PCPAPELLGGPSVFLFPPKPKDTLMISRTPEVTCVVVDVSHEDPEVKFNWYVDGVEVHNAKTKPREEQYNSTYRVVSVLTVLHQDWLNGKE<br>YKCKVSNKALPAPIEKTISKAKGQPREPQVYTLPPSRDELTKNQVSLTCLVKGFYPSDIAVEWESNGQPENNYKTTPPVLDSDGSFFLYSK<br>LTVDKSRWQQGNVFSCSVMHEALHNHYTQKSLSLSPGK |
| >PR1a_2G4_LC_CDS<br>ATGGGATTTGTTCTCTTTTCACAATTGCCTTCATTTCTTCTTGTCTCTACACTTCTCTTATTCCTAGTAATATCCCACTCTTGCCGTGCAG<br>GTGATATTCAAATGACTCAATCTCCAGCTTCTCTTTCTGTTTCTGTTGGAGAAACTGTTTCTATTACTTGTAGAGCTTCTGAAAATATTTA<br>TTCTTCTCTTGCTTGGTATCAACAAAAGCAAGGAAAGTCTCCACAACTTCTTGTTTATTCTGCTACTATTCTTGCTGATGGAGTTCCATCT<br>AGATTTTCTGGATCTGGATCTGGAACTCAATATTCTCTTAAGATTAATTCTCTTCAATCTGAAGATTTTGGAACTTATTATTGTCAACATT<br>TTTGGGGAACTCCATATACTTTTGGAGGAGGAACTAAGCTTGAAATTAAGAGGACTGTTGCTGCTCCATCTGTTTTCATCTTCCCACCATC<br>TGATGAGCAGCTCAAGTCTGGAACTGCTTCTGTTGTTTGCCTCCTCAACAATTTCTACCCAAGGGAAGCTAAGGTGCAGTGGAAAGTTGAT<br>AATGCTCTCCAGTCCGGAAACTCCCAAGAATCTGTTACTGAGCAGGATTCCAAGGATTCTACTTACTCCCTCTCCTCAACTCTCACTCTCT<br>CTAAGGCTGATTACGAGAAGCACAAGGTTTACGCTTGCGAAGTTACTCACCAGGGACTTTCTTCACCAGTGACTAAGTCTTTCAATAGGGG<br>CGAGTGCTAA |
| >PR1a_2G4_LC_aa<br>MGFVLFSQLPSFLLVSTLLLFLVISHSCRAGDIQMTQSPASLSVSVGETVSITCRASENIYSSLAWYQQKQGKSPQLLVYSATILADGVPS<br>RFSGSGSGTQYSLKINSLQSEDFGTYYCQHFWGTPYTFGGGTKLEIKRTVAAPSVFIFPPSDEQLKSGTASVVCLLNNFYPREAKVQWKVD<br>NALQSGNSQESVTEQDSKDSTYSLSSTLTLSKADYEKHKVYACEVTHQGLSSPVTKSFNRGEC |
| >PR1a_2G4_HC_CDS<br>ATGGGATTTGTTCTCTTTTCACAATTGCCTTCATTTCTTCTTGTCTCTACACTTCTCTTATTCCTAGTAATATCCCACTCTTGCCGTGCAG<br>GTGAAGTTCAACTTCAAGAATCTGGAGGAGGACTTATGCAACCAGGAGGATCTATGAAGCTTTCTTGTGTTGCTTCTGGATTTACTTTTTC<br>TAATTATTGGATGAATTGGGTTAGACAATCTCCAGAAAAGGGACTTGAATGGGTTGCTGAAATTAGACTTAAGTCTAATAATTATGCTACT<br>CATTATGCTGAATCTGTTAAGGGAAGATTTACTATTTCTAGAGATGATTCTAAGAGATCTGTTTATCTTCAAATGAATACTCTTAGAGCTG<br>AAGATACTGGAATTTATTATTGTACTAGAGGAAATGGAAATTATAGAGCTATGGATTATTGGGGACAAGGAACTTCTGTTACTGTTTCTTC<br>TACTTCAACTAAGGGACCATCTGTTTTTCCACTCGCTCCATCCTCTAAGTCTACTTCAGGTGGAACTGCTGCTCTTGGATGCCTTGTTAAG<br>GATTACTTTCCAGAGCCAGTGACTGTGTCTTGGAATTCTGGTGCTCTTACTTCCGGTGTTCATACTTTCCCAGCTGTGCTTCAATCTTCCG<br>GACTTTACTCTCTTTCCTCTGTTGTGACTGTGCCATCTTCTTCACTTGGCACTCAAACTTACATCTGCAACGTGAACCACAAGCCATCCAA<br>CACAAAAGTGGATAAGAAGGTTGAGCCAAAGTCCTGCGATAAGACTCATACTTGTCCACCATGTCCAGCTCCAGAACTTCTTGGTGGTCCT<br>TCTGTTTTTTTGTTCCCACCAAAGCCAAAGGATACTCTCATGATCTCTAGGACTCCAGAGGTTACATGCGTTGTGGTTGATGTGTCTCATG<br>AAGATCCAGAGGTGAAGTTCAACTGGTATGTGGATGGTGTTGAGGTGCACAACGCTAAGACTAAGCCAAGAGAGGAACAGTACAACTCCAC<br>TTACAGGGTTGTGTCTGTGCTTACTGTTCTTCACCAGGATTGGCTTAACGGCAAAGAGTACAAGTGCAAGGTGTCCAACAAGGCTTTGCCA<br>GCTCCAATCGAAAAGACTATCTCTAAGGCTAAGGGACAGCCAAGGGAACCTCAAGTTTACACTCTTCCACCATCTAGGGATGAGCTTACTA<br>AGAACCAGGTGTCCCTTACTTGCCTTGTGAAGGGATTTTACCCATCCGATATTGCTGTTGAGTGGGAGTCTAATGGACAGCCTGAGAACAA |

| |
|---|
| **CTACAAGACTACTCCACCAGTGCTCGATTCCGATGGATCATTCTTCTTGTACTCCAAGCTCACTGTGGATAAGTCTAGGTGGCAACAGGGA<br>AACGTTTTCTCTTGCTCTGTTATGCATGAGGCTCTCCACAATCACTACACTCAGAAGTCCCTTTCTTTGTCCCCTGGCAAGTAA** |
| >PR1a_2G4_HC_aa<br>**MGFVLFSQLPSFLLVSTLLLFLVISHSCRAGEVQLQESGGGLMQPGGSMKLSCVASGFTFSNYWMNWVRQSPEKGLEWVAEIRLKSNNYAT<br>HYAESVKGRFTISRDDSKRSVYLQMNTLRAEDTGIYYCTRGNGNYRAMDYWGQGTSVTVSSTSTKGPSVFPLAPSSKSTSGGTAALGCLVK<br>DYFPEPVTVSWNSGALTSGVHTFPAVLQSSGLYSLSSVVTVPSSSLGTQTYICNVNHKPSNTKVDKKVEPKSCDKTHTCPPCPAPELLGGP<br>SVFLFPPKPKDTLMISRTPEVTCVVVDVSHEDPEVKFNWYVDGVEVHNAKTKPREEQYNSTYRVVSVLTVLHQDWLNGKEYKCKVSNKALP<br>APIEKTISKAKGQPREPQVYTLPPSRDELTKNQVSLTCLVKGFYPSDIAVEWESNGQPENNYKTTPPVLDSDGSFFLYSKLTVDKSRWQQG<br>NVFSCSVMHEALHNHYTQKSLSLSPGK** |

The N-terminal signal peptide of *Nicotiana tabacum* pathogenesis-related protein PR1a (grey) was used, followed by a glycine linker (orange), the LC/HC of 2F5 and a stop codon (red).

**Table S2** Plasmids used in this study

| Plasmid | Alternative name | Description | Reference |
|---|---|---|---|
| pJK268c | 35SP_P19_35ST | p19 RNA silencing suppressor | Kourelis et al., 2020 |
| pKB002 | EV_control | Empty vector control | This work |
| pKB018 | L0_2F5_LC | LC of 2F5 (CDS) | This work |
| pKB019 | L0_2F5_HC | HC of 2F5 (CDS) | This work |
| pKB020 | L1_35SP_2F5_LC_nosT | LC of 2F5 (TU) | This work |
| pKB021 | L1_35SP_2F5_HC_nosT | HC of 2F5 (TU) | This work |
| pKB024 | L0_2G4_LC | LC of 2G4 (CDS) | This work |
| pKB025 | L0_2G4_HC | HC of 2G4 (CDS) | This work |
| pKB026 | L1_35SP_2G4_LC_nosT | LC of 2G4 (TU) | This work |
| pKB027 | L1_35SP_2G4_HC_nosT | HC of HC (TU) | This work |
| pICH41308 | L0_CDS | Golden Gate Level 0 acceptor for CDS | Engler et al., 2014 |
| pICH51277 | L0_35SP_5UTR | Golden Gate Level 0 35S promoter | Engler et al., 2014 |
| pICH41421 | L0_nosT | Golden Gate Level 0 nos terminator | Engler et al., 2014 |
| pICH47732 | L1_P1_Fwd | Golden Gate Level 1 Position 1 Forward acceptor | Engler et al., 2014 |
| pICH47732 | L1_P2_Fwd | Golden Gate Level 1 Position 2 Forward acceptor | Engler et al., 2014 |
| pFGH008 | L1_35SP_NbPR4_35ST | Cys Inhibitor | Grosse-Holz et al., 2018b |
| pFGH010 | L1_35SP_SICDI_35ST | Ser/Asp Inhibitor | Grosse-Holz et al., 2018b |
| pFGH047 | L1_35SP_HsTIMP_35ST | Metalloprotease Inhibitor | Grosse-Holz et al., 2018b |
| pFGH048 | L1_35SP_EPI1_35ST | Ser/Cys Inhibitor | Grosse-Holz et al., 2018b |
| pFGH049 | L1_35SP_EPI12_35ST | Ser/Cys Inhibitor | Grosse-Holz et al., 2018b |
| pFGH053 | L1_35SP_NbPot1_35ST | Ser Inhibitor | Grosse-Holz et al., 2018b |
| pFGH054 | L1_35SP_SICYS8_35ST | Cys Inhibitor | Grosse-Holz et al., 2018b |
| pFGH203 | L1_35SP_Ala-HsTIMP_35ST | HsTIMP inactive mutant | Grosse-Holz et al., 2018b |
| pFGH214 | L1_35SP_SICYS8-Q47P_35ST | SICYS8 inactive mutant | Grosse-Holz et al., 2018b |
| TRV1 | TRV1 | Tobacco Rattle Virus RNA (TRV) 1 | Liu et al., 2002 |
| pJK037 | TRV2gg (pL2M-TRV2) | TRV2 Golden Gate acceptor vector | Morimoro et al., 2022 |
| TRV2::PDS | TRV2::PDS | TRV2 for phytoene desaturase (PDS) gene | Liu et al., 2002 |
| pPB035 | TRV2::GFP | TRV2 for GFP | This work |
| pPB058 | TRV2::SBT1.7a | TRV2 for *sbt1.7a* | This work |
| pPB059 | TRV2::SBT1.7c | TRV2 for *sbt1.7c* | This work |
| pPB065 | TRV2::SBT1.9a | TRV2 for *sbt1.9a* | This work |
| pPB039 | TRV2::SBT5.2 | TRV2 for *sbt5.2a* | This work |
| pPJ048 | 35S_cytosolicGFP_35ST | Cytosolic GFP | Jutras et al., 2021 |
| pPJ057 | 35S_secretedRFP_35ST | Secreted RFP | This work |

CDS, coding sequence; TU, transcription unit; TRV, Tobacco Rattle Virus.

**Table S3** Virus-induced gene silencing (VIGS) fragments used in this study.

| **Name** | **Accession number (LAB360)** | **Sequence (5' - 3')** |
|---|---|---|
| *TRV::GFP* | N/A | ATGCGTAAAGGCGAAGAGCTGTTCACTGGTGTCGTCCCT<br>ATTCTGGTGGAACTGGATGGTGATGTCAACGGTCATAAG<br>TTTTCCGTGCGTGGCGAGGGTGAAGGTGACGCAACTAAT<br>GGTAAACTGACGCTGAAGTTCATCTGTACTACTGGTAAAC<br>TGCCGGTACCTTGGCCGACTCTGGTAACGACGCTGACTT<br>ATGGTGTTCAGTGCTTTGCTCGTTATCCGGACCATATGAA<br>GCAGCATGACTTCTTCAAGTCCGCCATGCCGGAAGGCTA<br>TGTGCAGGAACGCACGATTTCCTTT |
| *TRV::SBT1.7a* | NbLab360C02+56673784 | GCCGGACACGTGGATCCCGTTTCAGCATTAAACCCAGGA<br>CTCGTCTACGACATAACCGCCGACGATTATCTCAATTTCCT<br>GTGTGCATTGGATTACACTCCCTCACAAATCAGCAGCTTA<br>GCAAGAAGAAACTTCACGTGCAATGAGAGTAAAAAATACA<br>GTGTCACAGATTTGAATTACCCTTCGTTTGCAGTCTCATTT<br>CCAGCAGAATCCTCAGCTCGAACCGGGAGTGCCGGTTCA<br>AGCTCAATCAAATACAGCCGGACGCTGACTAATGTTGGAC<br>CGGCTGGAACATATAAAGTT |
| *TRV::SBT1.7c* | NbLab360C06+107534609 | TTGGTCTCAATTCCCAAAAAAAATGTCAAGATTTACAATGC<br>TAGTAGTTCTTGTTCTTGTTCTTCTATGTCTATGCCATTTAT<br>CAGTAGCAACAATAGGAAGTAGTAATAAGAAGAGTACTTAC<br>ATAGTACACGTGTCAAAATCCCAAATGCCAGAGAGTTTTG<br>AAGACCATAAACGCTGGTATGATTCATCACTAAAATCAGTT<br>TCTGATTCAGCAGAAATGTTGTATGTTTACAACAACGTTGT<br>ACATGGTTTCTCAGCAAGACTGACTGTTCAAGAAGCAGAA<br>TCACTTGAGAGACAATCTGGGATTCTGGGATTGAGACCAA |
| *TRV::SBT1.9a* | NbLab360C18-103095902 | ATGGCAAATCATATTACCTTGTGTATTTGGTTGCTTTTCTTC<br>TTTATTTCTATATTTTCACTAGCAAAGTCAGAAACATATATCA<br>TTCATATGGATTTGTCAGCCATGCCAAAAGCTTTTTCTAGC<br>CATCATAATTGGTACTTGACAACACTTTCTTCTGTATCAGA<br>CAGCAGTACAAATCACAAAGACTTCTTGTCCTCAAAACTA<br>GTCTATGCTTATACTAATGCTATACATGGTTTTAGTGCAAGT<br>CTCTCTCCTTCTGAACTTGAAGCCATAAAATATTCTCCAGG<br>TTATGTTTCT |
| *TRV::SBT5.2* | NbLab360C13-25441362 | TATGAATTATCCATCAATAGCTATTTCTAGTCTCAAAGAAAA<br>CGAGCCGAAGAAAGTTACTAGAACTGTAACAAATACTGGT<br>GAAGAAGCATCAGTATATACTACAGTTATTGAGGCACCAAA<br>AGGATTGGAAGTCCAAGTGATCCCAACTAAATTGGAATTTA<br>CAAATAAAAGCAAGAAATTAAGCTATGATGTGTCTTTCAAA<br>GCTTCATCTACCTCAAAGGAAGATCTGTTTGGATCAATTAC<br>TTGGACTAATGGTAAGTACAAAGTCCGGAGTCCATTCGTC<br>GTAAGTAGCAACTA |

**Table S4** Single-guide RNAs (sgRNAs) and primers used for sequencing of *sbt5.2* KO lines.

| **Gene** | **sgRNAs** | **Sequencing primers (5' - 3')** |
|---|---|---|
| *sbt5.2a* | GCCAGGGACGAGAACGGACA | F:GTGACTGCGTCATTGATGATTAG<br>R:CTATGGCAATAGGATTGCTTGG |
| | GGGGTGGATCTCCTGGTTCG | |
| *sbt5.2b* | GGCGCGTTTATCAGAAGTTG | F:GTGGTTCTCCAGGATCATATCAA<br>R:CAAGATTCCGATTATGGTGTCAG |
| | GCTGCAAAGGGTGGGTCGCC | F:AGATTTTATGATGACTCTGCTGA<br>R:TTGTAGTAGCTCCCACTGTGAGA |
| *sbt5.2c* | GCCATGTCCATTCTTGTCCC | F:AGTCCTATCTCGTTTTGATTATT<br>R:CTATTTCCAGCAGAACAGACAAC |
| | GGCACATCTAATGGATGCCG | |
